# Phase precession in the human hippocampus and entorhinal cortex

**DOI:** 10.1101/2020.09.06.285320

**Authors:** Salman E. Qasim, Itzhak Fried, Joshua Jacobs

**Author notes:** Address correspondence to: Joshua Jacobs, 351 Engineering Terrace, Mail Code 8904, 1210 Amsterdam Avenue, New York, NY 10027.

## Abstract

Knowing where we are, where we have been, and where we are going is critical to many behaviors, including navigation and memory. One potential neuronal mechanism underlying this ability is phase precession, in which spatially tuned neurons represent sequences of positions by activating at progressively earlier phases of local network theta (~5–10 Hz) oscillations. Phase precession may be a general neural pattern for representing sequential events for learning and memory. However, phase precession has never been observed in humans. By recording human single-neuron activity during spatial navigation, we show that spatially tuned neurons in the human hippocampus and entorhinal cortex exhibit phase precession. Furthermore, beyond the neural representation of locations, we show evidence for phase precession related to specific goal-states. Our findings thus extend theta phase precession to humans and suggest that this phenomenon has a broad functional role for the neural representation of both spatial and non-spatial information.

## Introduction

Our brain’s ability to link related experiences is critical to everyday life, and to our memory for past experiences. One crucial example is spatial navigation, which requires awareness of individual locations and the association between multiple locations, such as those on the same path. Similarly, episodic memory requires the encoding of individual events and the association between sequential events. The hippocampal formation is important for both spatial cognition and episodic memory^1–4^. Therefore, neural activity in this region might play a key role in linking sequential locations and events.

Specifically, theories of how the brain represents sequential experiences rely on the idea that the timing of neuronal spiking is critical for learning associations^5–10^. Spike timing, in turn, is thought to be coordinated across networks by fluctuations in the large-scale network activity that can be estimated via the local field potential (LFP)^11–16^. This suggests that a coordinated relationship between network oscillations and single-cell spiking may play a mechanistic role in complex behaviors or aspects of cognition, such as memory^17,18^, that require the association of sequential events. A prominent potential mechanism for the binding and compressing of sequential events is hippocampal phase precession, extensively observed in rodents, during which active neurons rhythmically spike in coordination with the local theta frequency (5-10 Hz) oscillation. Phase precession is readily observable in many hippocampal place cells^19^ and entorhinal grid cells^20,21^. Because these neurons spike slightly faster than the theta oscillation as the rodent runs through specific locations, phase precession results in sequences of locations being encoded at different phases of theta oscillations. As such, phase precession may compress spatial trajectories on the scale of behavior into the brief timescale of the theta cycle that is conducive to synaptic plasticity^22–24^.

There is evidence that phase precession’s utility for binding and compressing sequential events may be used by the brain to represent non-spatial features of experience as well. While phase precession is often described in hippocampal place cells or entorhinal grid cells, it has also been observed in a diverse range of brain areas such as ventral striatum^25^, subiculum^26^, basal forebrain^27^, and medial prefrontal cortex^28^. Critically, a slew of recent work has directly observed phase precession independent of location within a place or grid field, encoding elapsed time during REM sleep^29^, wheel-running^30^, jumping^31^, fixation^32^, presentation of task-relevant stimuli^33–35^, and task epoch^27^. The widespread prevalence of phase precession suggests that this phenomenon has a more general role beyond representing the current spatial location, and that it could be relevant for building neural representations in many regions to support diverse aspects of cognition, learning, and memory.

Despite its prevalence in rodents, and the many theories suggesting a fundamental role for phase precession in neural coding^19,36–38^, phase precession has not been observed in humans. To examine this issue, we analyzed simultaneous single-neuron and LFP activity from neurosurgical patients as they performed a virtual spatial memory task and examined the patterns of spike–LFP interaction. Here, we describe spatial phase precession in humans analogous to that observed in navigating rodents. We also describe evidence for phase precession related to the coding of non-spatial variables, in which neurons transiently spike faster than the theta oscillation during trajectories to specific goals. Overall, our findings extend precession to humans and demonstrate its potential use for encoding both spatial and non-spatial features of experience.

## Results

### Spatial phase precession in hippocampus and entorhinal cortex during navigation

We analyzed recordings of neuronal spiking from 1,074 neurons in the hippocampus, entorhinal cortex, parahippocampal gyrus, anterior cingulate cortex, and amygdala of 13 neurosurgical patients undergoing clinical treatment for drug-resistant epilepsy. During recordings, subjects performed a goal-directed navigation task in a 2D virtual environment on a laptop computer^39,40^ (Supplementary Fig. 1; see *Methods*). The virtual environment contained six goal stores surrounding the perimeter of a square track, with the center of the environment obstructed by buildings. Subjects were able to travel around the track in either clockwise or counterclockwise directions (Fig. 1A).

**Figure 1:**
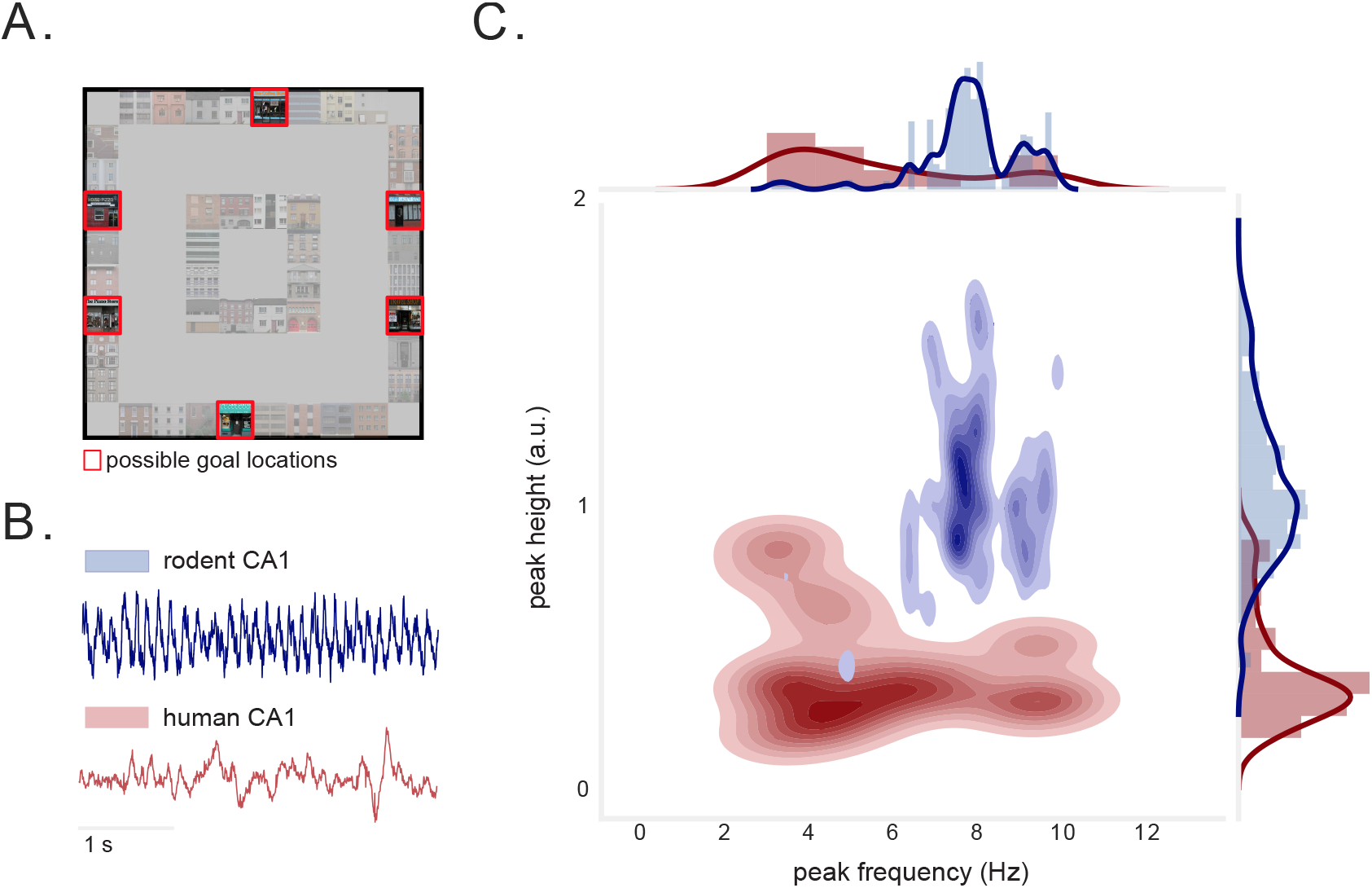
Virtual environment and hippocampal theta oscillations during task. A) Overhead view of task environment. Red squares denote locations of possible goal locations. B) Examples of raw LFP data from rodent (publicly available dataset^41^) and human hippocampus. C) Joint distribution of peak frequency and peak height of LFP power spectral density (PSD) from rodent (blue) and human (red) hippocampus. Rodent hippocampal recordings exhibit highly stereotyped peaks. Human hippocampal recordings exhibit significantly smaller peak heights, and peaks are at significantly lower frequencies (*p*’s< 2 × 10^-34^, Wilcoxon rank-sum tests).

Given our interest in phase coding, we first characterized the prevalence of theta oscillations in the human hippocampus and compared their properties to those seen in rodents, leveraging a publicly-available dataset^42^.

Compared to rodents, human hippocampal theta spanned a significantly broader range of frequencies (*p* < 4 × 10^-4^, Levene test), with significantly smaller, lower-frequency peaks in the power spectrum (*p*’s< 3 × 10^-8^, Wilcoxon rank-sum tests; Figs. 1B-C, S2). Because human theta appears to span both low and high frequencies^43^, we assessed phase precession with respect to oscillations at a broad range of LFP theta frequencies (2–10 Hz) (Fig. 1C). To assess phase precession we first identified each neuron whose firing was modulated by the subject’s position in the virtual environment. We labelled the clockwise (CW) and counter-clockwise (CCW) movement periods and then used a shuffle-corrected ANOVA to identify 292 spatially modulated neurons that fired significantly more when subjects moved through particular locations during one or both of these movement conditions^44^, after correcting for multiple comparisons (see *Methods*). Because phase precession in rodents is most predominant near the place-field center^45^ and on short trajectories^46^, we tested for phase precession during short trajectories through the field center, defined as the peak firing location for each neuron (Fig. 2A).

**Figure 2:**
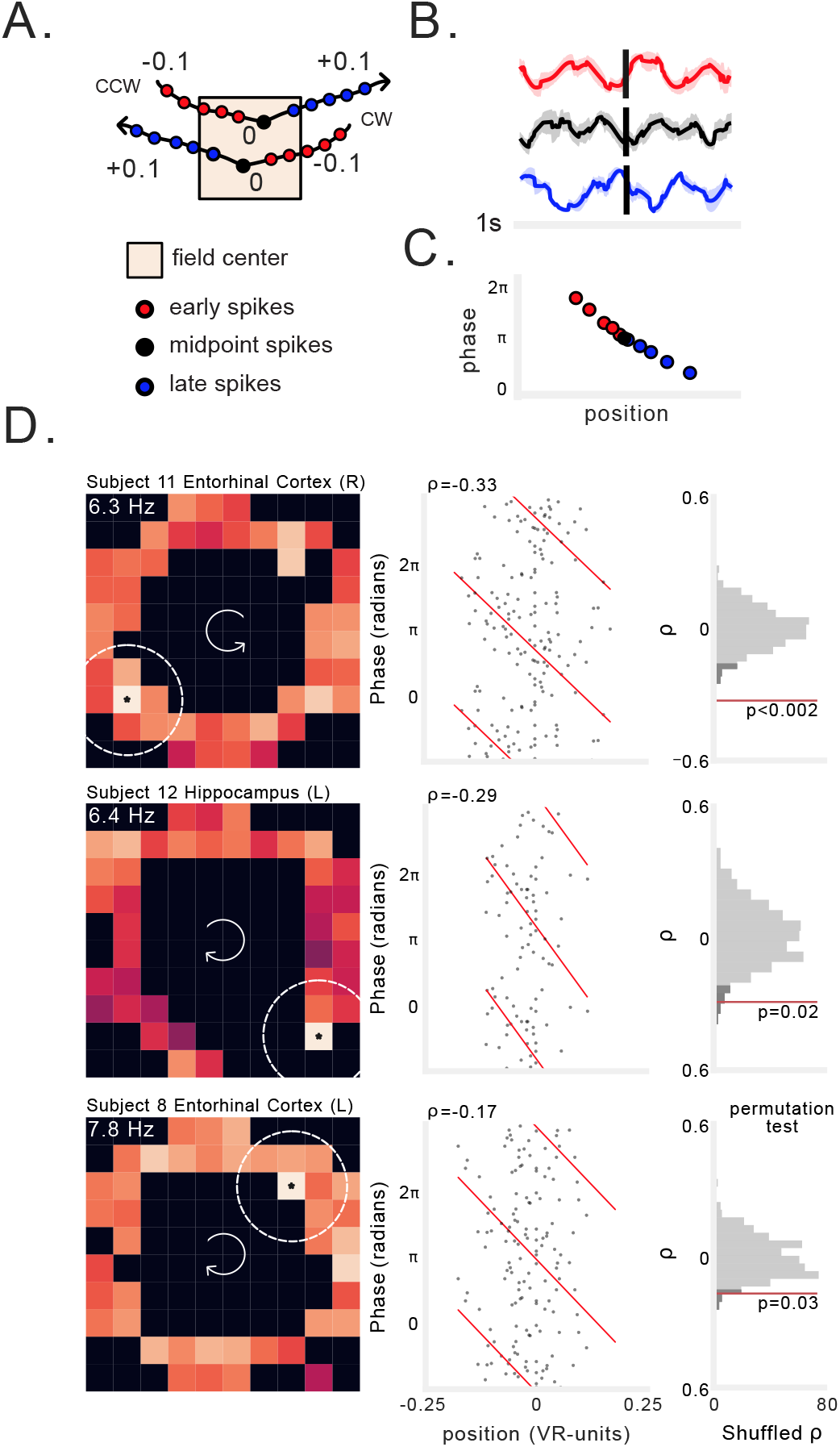
Examples of spatial phase precession in human hippocampus and entorhinal cortex. A) Schematic illustrating our method for selecting spikes near peak firing bin (*see Methods*). For each spatially modulated cell we analyzed phase precession using spikes that occurred early (red), at the midpoint (black), and late (blue) in trajectories through the center of the firing field. B) Spike-triggered average (STA) LFP (reconstructed from phase) for early, midpoint, and late trajectory spikes from one neuron. C) Schematic of spike phase as a function of distance from center spike during a trajectory through the field, showing phase precession as a negative progression of phase-by-position. D) Three examples of spatial phase precession. Each row shows an individual neuron. Left: firing rate heat map. Text label indicates average firing rate in peak firing bin, which is noted with an asterisk. Brighter colors denote higher firing rates. Dotted lines indicate maximum radius around field in which spiking was assessed. Arrows in the center of the heatmap indicate the movement direction. Middle: spike phase as a function of location relative to the field center. Spike phases are duplicated vertically to enable visualization of circular-linear regression (red). Rho indicates circular-linear regression coefficient. Right: statistical assessment of circular-linear regression rho using surrogate distribution of circular-linear regression rho values generated by permutation of spike phases. Red line indicates value of real data. Dark gray shading indicates 95^*th*^ percentile of surrogate distribution.

We observed that some of the spatially tuned neurons showed spiking at progressively earlier phases of the theta oscillation during individual trajectories through their firing field (Supplementary Fig. 3). To assess if this was a consistent pattern across trajectories, we leveraged the fact that during phase precession, spikes at later positions in the trajectory should occur at earlier phases, manifesting as a negative correlation between spike-phase and position^19^. In this way, spiking at different phases of the LFP would correspond to different relative positions along the path to a neuron’s firing field center (Fig. 2B,C). We tested for this pattern by measuring the correlation^47^ between spike-phase and position using circular statistics^48^ and a shuffle-based permutation procedure (see *Methods*).

By performing this procedure across all of the spatially-tuned neurons we identified, we report the first evidence of phase precession in humans. Figure 2D shows three examples of single neurons recorded in the hippocampus and entorhinal cortex that exhibited significant phase precession during navigation at particular spatial locations (see Supplementary Fig. 4 for additional examples). Each of these neurons increased their firing in a specific region of the environment (Fig. 2D, left). As a person approached the center of that region, the neuron initially spiked at late phases of the 2–10 Hz LFP, but as they continued their trajectory through the center and past it, spikes occurred at progressively earlier phases (Fig. 2D, middle). This change in spike phases between early positions and late positions is characterized by significant negative phase–position correlations (Fig. 2D, right).

After testing the spatially-tuned neurons in our dataset for phase precession with our permutation procedure, we found that precession was widespread, with 12% (35/292) of neurons exhibiting this phenomenon, which is well above what would be observed by chance (*p* < 3 × 10^-6^, binomial test; Fig. 3A). Of these 35 neurons, 22 neurons exhibited uni-directional spatial tuning and precession and 10 neurons exhibited bi-directional spatial tuning and precession. The remaining 3 neurons exhibited uni-directional precession in one location and bi-directional precession in another. Notably, we specifically observed significant proportions of spatially modulated cells exhibiting spatial phase precession in the hippocampus and entorhinal cortex (*p*’s< 0.002, binomial test; Fig. 3B). Phase precessing neurons exhibited an average circular–linear correlation coefficient of −0.26 ± 0.09, an average slope of −1.36 ± 0.8 radians/VR-units (Fig. 3C), and an average in-field firing rate of 4.9 ± 1.7 Hz (Fig. 3D). Phase precessing neurons had spatial firing fields throughout the environment (Fig. 3E). To test whether spatial phase precession was consistent across an entire behavioral session, we separately computed the circular–linear correlation coefficient for the first and second halves of the session and found no difference between halves (*p* = 0.4, paired t-test), with significantly negative correlation coefficients in each half (*p*s< 0.002, one-sample t-test). We found evidence for spatial phase precession in 11/12 of the subjects with spatially modulated neurons. These results thus demonstrate the existence of phase precession as a neural code for spatial position in humans during virtual navigation. The theta-frequency (2–10 Hz) and regions involved (hippocampus, entorhinal cortex) suggest that the phase precession we observed is largely analogous to that documented in rodent place and grid cells.

**Figure 3:**
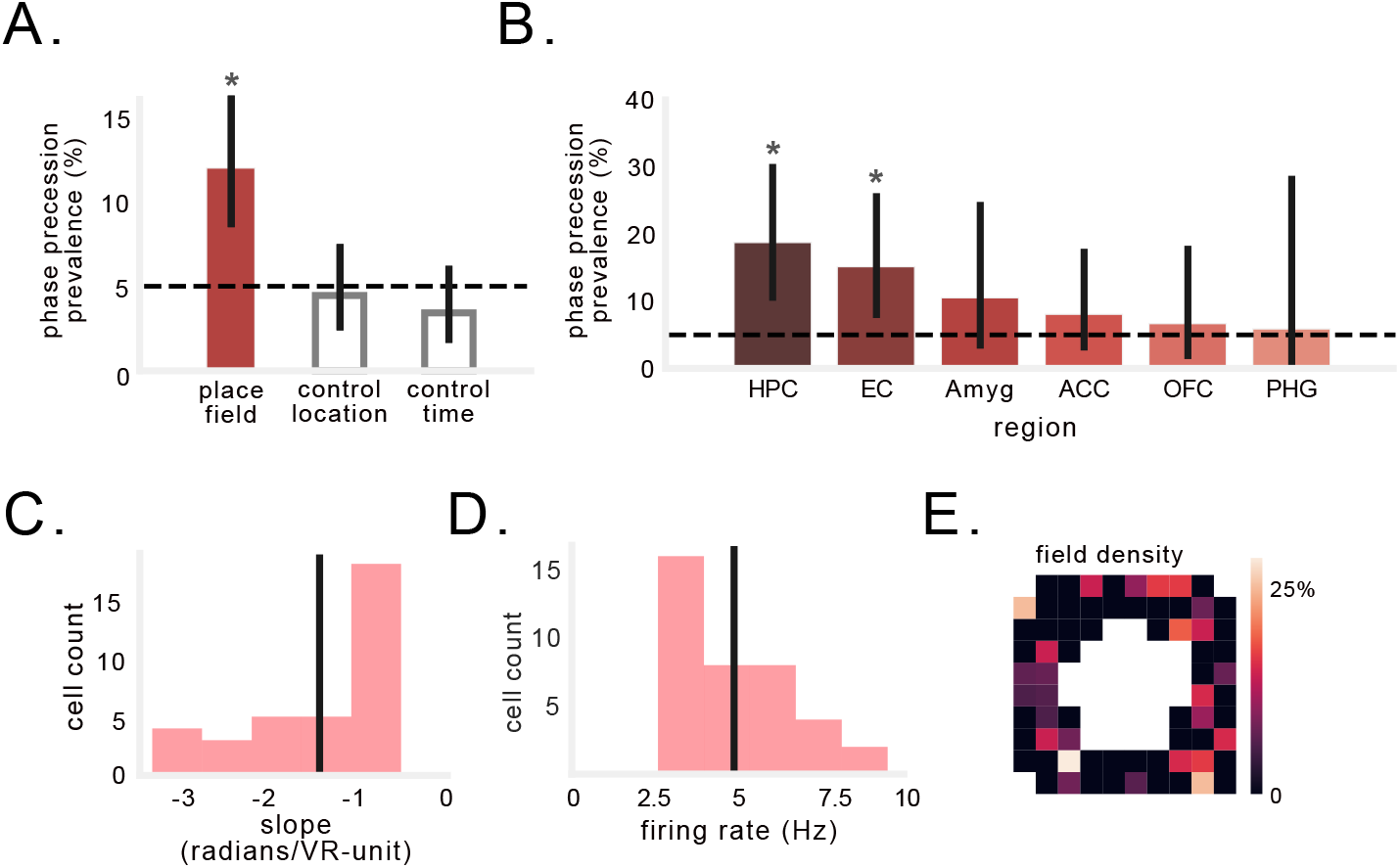
Prevalence and characteristics of spatial phase precession in humans. A) Percentage of spatially modulated cells that exhibit phase precession during trajectories through the firing field (filled bar). Grey bars show control analyses of precession relative to alternative locations, or as a function of time, not position, during spiking episodes (see Supplementary Fig. 5). Black dotted line denotes chance. Solid black line indicates 95% binomial confidence interval. Asterisk indicates significant proportion of spatially modulated cells exhibiting phase precession during trajectories through the firing field (*p* < 3 × 10^-6^, binomial test). B) Percentage of spatially modulated cells across regions. Asterisk indicates significant proportion of cells exhibit phase precession (*p*s< 0.002, binomial test). C) Distribution of circular-linear regression slopes identified for neurons exhibiting significant phase precession. Black line indicates mean. D) Distribution of average firing rate of peak firing bins in which phase precession was observed. E) Prevalence of phase precession across the environment. Colors indicate percentage of firing fields in each bin that exhibited precession.

To be sure that our findings of precession indicated a spatial phase code relative to space, we tested two alternative explanations for our results: that precession was equally prevalent at randomly selected spatial locations (in which the neuron was sufficiently active), or that precession was actually measuring the advance of spike phase according to elapsed time (see *Methods*; Supplementary Fig. 5A, C). Neither alternative model identified significant proportions of phase-precessing cells, and these models resulted in the identification of a smaller number of cells as compared to our primary analyses (*χ*^2^ = 20.6, *p* < 4 × 10^-5^, chi-squared test; Fig. 3A; see also Supplementary Fig. 5B, D). These results indicate that human phase precession occurs more strongly at locations that show the highest firing rates, and that phase precession in spatially tuned neurons is more closely tied to location than elapsed time during movement.

### Evidence for phase precession without spatial coding

While phase precession has been observed most readily relative to specific spatial locations, there is also evidence for precession with respect to non-spatial behaviors and stimuli^27,29–31,33^, and in regions outside the hippocampal formation^25,28^. These findings suggest that phase precession could be a more general phenomenon that the brain uses to represent diverse types of consecutive, relevant stimuli/states using different phases of an oscillation. To examine this possibility, we used a broader analytical method to test whether the non-spatially tuned neurons exhibited phase precession without reference to position. To do this, we measured each neuron’s rhythmic frequency of spiking in comparison to the local theta oscillation^49,50^. Identifying a consistent pattern of faster-than-LFP rhythmic spiking would indicate the presence of a precession-like pattern of LFP-coordinated spiking that could bind and compress sequential, non-spatial features of the task — just as spatial phase precession is theorized to do for locations^24,51^.

We identified neurons that showed rhythmic spiking at a frequency faster than the theta oscillation by using a method that has identified this pattern in animals with very stereotyped, 8 Hz theta such as rats^26,42,52^ and mice^53,54^, as well as animals with human-like theta that appears at a range of frequencies, such as bats^55^. In brief, in this method we first measured the theta phase estimate for each spike from the concurrent 2–10 Hz LFP and “unwrapped” the phase time series so that it increased linearly, like elapsed time. We then measured the spike-phase spectrum, which we defined as the power spectral density of the time series of unwrapped spike phases (*see Methods*). In contrast to conventional spectral analysis that measures the frequency of a signal relative to absolute time, the spike–phase spectra reflects the relative frequency of rhythmic spiking compared to the frequency of concurrent LFP oscillations. If a spike-phase spectra showed a peak at a relative frequency > 1.0 it would indicate that a neuron’s rhythmic spiking was faster than the concurrent oscillations in the LFP, and thus this neuron’s spiking exhibited precession relative to the LFP (Fig. 4A). This method ensures that a consistent relationship between the spiking frequency and LFP frequency can be identified even if the LFP shifts in frequency or amplitude, and even though neuronal spike times alone may not show a clear oscillation (Supplementary Fig. 6), as is the case in humans and bats^55^. We validated this method by applying it to data from rodent CA1 and identifying a consistent > 1.0 relative frequency (Supplementary Fig. 7), consistent with the spatial phase precession observed in these neurons^42^.

**Figure 4:**
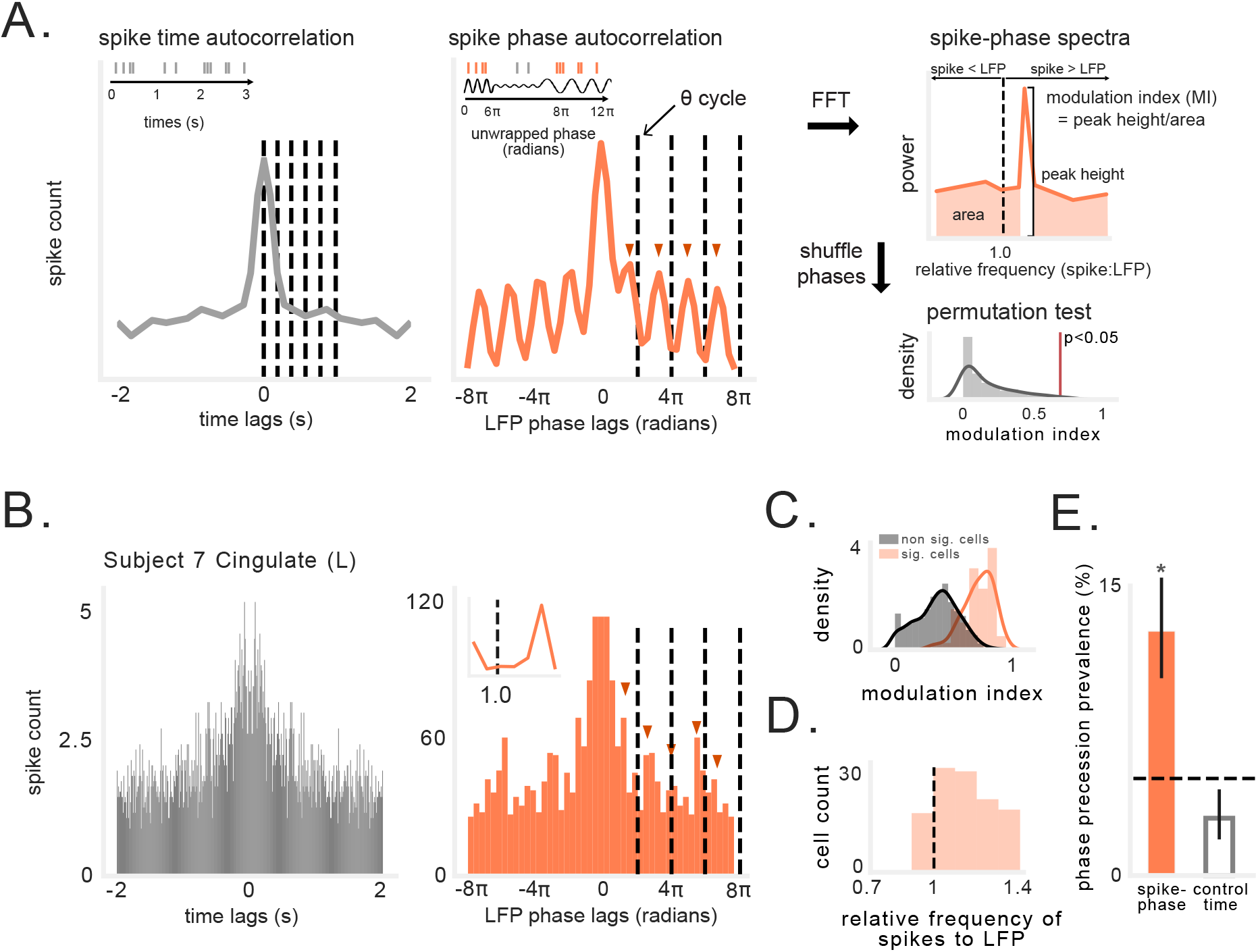
Spike-phase spectra reveals precession-like pattern in non-spatially tuned neurons. A) Schematic illustrating analysis of rhythmic spiking frequency relative to LFP oscillation (*see Methods*). Left: Autocorrelation of spike times (gray), with dotted lines at 200 ms intervals. Middle: Autocorrelation of unwrapped spike phases. Dotted lines indicate one cycle of ongoing LFP in 2–10 Hz band. Red arrows indicate peaks in autocorrelation, which occur progressively earlier than cycles of ongoing LFP. Fast Right: Fourier transform (FFT) of autocorrelation function yields power spectral density(PSD) showing cell spiking frequency relative to ongoing LFP frequency. The modulation index (MI) is visualized here as the ratio of the spectral peak height to power at all other relative frequencies. This value is compared to a null distribution of MI values generated by shuffling spike phases in each cycle. B) Left: Spike time autocorrelation showing little evidence of theta modulation. Right: Spike phase autocorrelation (orange) showing cell oscillating slightly faster than ongoing LFP (cycles of 2–10 Hz LFP indicated by dotted line). Inset shows spike-phase spectra. C) Modulation index (MI) of spike-phase spectral peaks for significant vs. non-significant neurons. D) Distribution of relative frequencies for neurons exhibiting significant MI. Values to the right of the black line indicate that the rhythmic spiking frequency slightly exceeded the LFP frequency. E) Percentage of non-spatial cells that exhibit precession-like spiking relative to LFP phase, compared to cell’s exhibiting precession relative to time in a spiking episode. Black dotted line denotes chance level. Solid black line indicates 95% binomial confidence interval. Asterisk indicates significant proportion of cells (*p* < 7 × 10^-18^, binomial test).

To assess whether precession-like rhythmic spiking was evident for non-spatially tuned cells, we applied this method to the 744 neurons that were active during the task but did not exhibit significant spatial tuning. Figure 4B depicts an example neuron that we identified with this method that showed significant precession. This analysis found that the rhythmicity of this cell’s spiking frequency occurred at a frequency that reliably exceeded the frequency of the LFP (right panel), although no consistent rhythm is apparent from the spike timing alone (left panel). Using this method we found that 12% of non-spatially tuned neurons (90/744) showed a significant relationship between neuronal spiking frequency and LFP frequency (Fig. 4C), at a range of relative frequencies > 1.0 (Fig. 4D). The number of neurons that showed this phenomenon was significantly more than we expected by chance (*p* < 7 × 10^-18^, binomial test; Fig. 4E).

We performed a control analysis (Supplementary Fig. 5C) to rule out the possibility that these effects could be explained by the absolute spike timing, though this was unlikely given the relative lack of intrinsic rhythmicity in the spike timing (see Supplementary Fig. 6). This analysis confirmed that most of these neurons show phase precession only when spiking is measured relative to the instantaneous ongoing oscillation rather than absolute time^56^ (Fig. 4E). These results illustrate how the frequency variability of human hippocampal theta^43^ may diminish traditional measures of phase precession, and demonstrate the potential for phase precession in neurons that are not spatially tuned. We next sought to test whether this new non-spatial precession pattern might vary behaviorally in relation to non-spatial, higher level features of the task, such as prospective goals.

### Evidence for phase precession during trajectories to specific goals

Having shown that non-spatially tuned neurons can exhibit phase precession, we next tested whether this was a tonic pattern^50^ or, alternatively, one that emerged selectively to code for specific stimuli or states. Specifically, recent work has shown that human hippocampal– cortical networks represent goals and their intermediate locations^57^; furthermore we found previously that this task elicits distinctive patterns of rate- and phase-coding for goals^58^. Therefore, we assessed whether phase precession emerged selectively during trajectories to specific goals in service of binding those trajectories for learning and memory.

During each trial of this task, the subject was cued to travel to a randomly selected goal location (Fig. 5A). We found that some neurons specifically showed phase precession only during travel to particular goals. Figure 5B shows an example of a neuron whose spiking shows phase precession during navigation to goal 2, but not the other goals. This effect is evident in the spike-phase autocorrelogram for that goal, which shows that during travel to goal 2 rhythmic spiking occurs at a frequency that is slightly faster than the ongoing 2–10 Hz LFP. To systematically test for goal-state phase precession, we measured the spike-phase spectrum during trajectories to each goal and compared these spectra between goals, using a permutation procedure and correcting for multiple comparison across goals (Fig. 5C, see *Methods*). Figure 5D depicts two example neurons from the hippocampus and amygdala of two different subjects (see Supplementary Fig. 8 for additional examples). These neurons exhibited rhythmic spiking at faster frequencies than the ongoing LFP while the subjects were en route to specific goals. Critically, this rhythmic spiking was goal-specific and did not appear during trajectories to other goals. These patterns were thus examples of phase precession for a particular goal-state, similar to phase precession in a place field.

**Figure 5:**
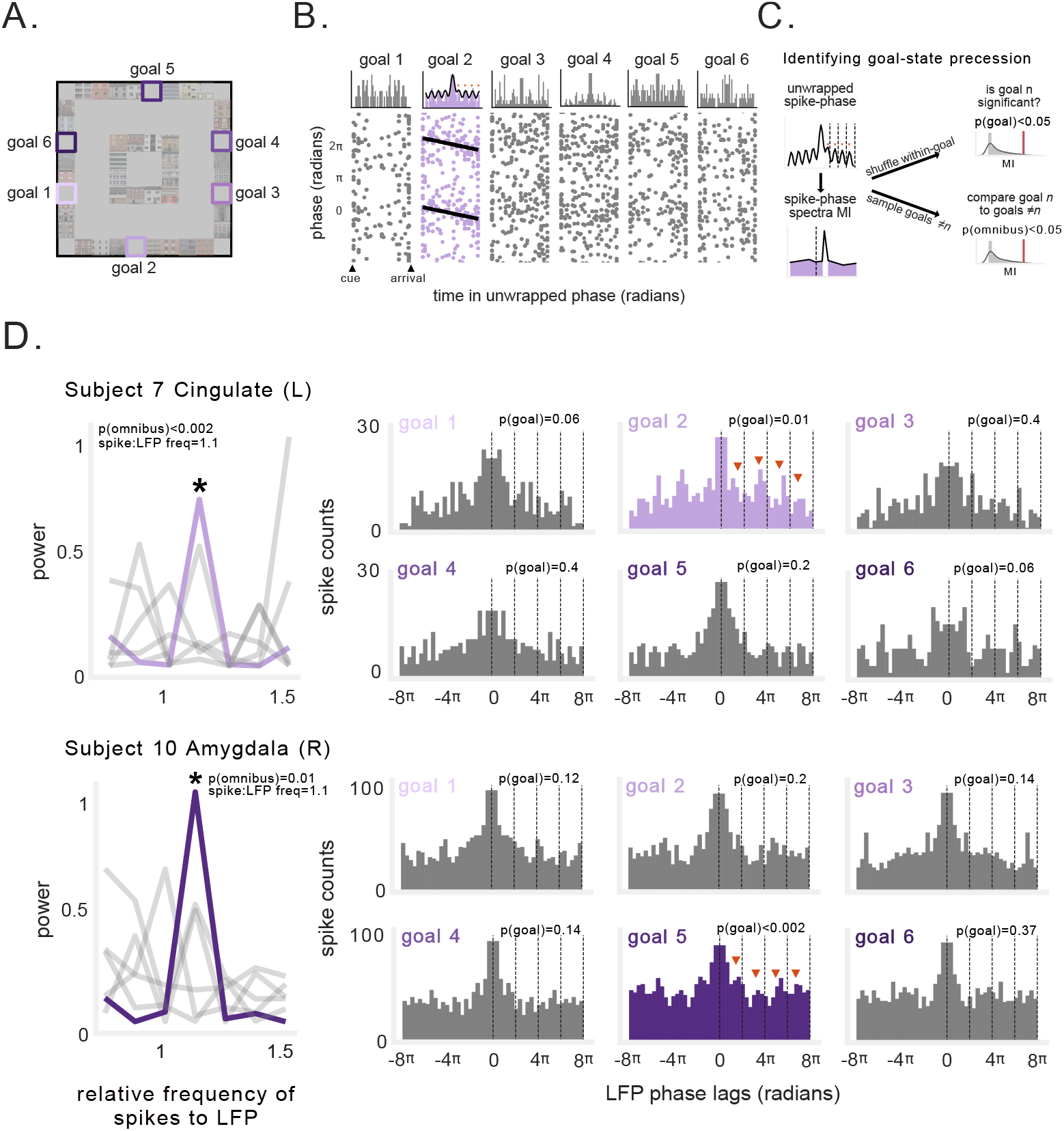
Goal-state phase precession. A) Schematic of task environment. Labels indicate goal locations. B) Spike-phases during navigation to different goals for example neuron. Top: unwrapped spike-phase autocorrelograms for each goal. Black line indicates fit of decaying-oscillation function. Spiking frequency transiently exceeded LFP frequency only during navigation to goal 2. Bottom: Spike-phase as a function of duration within each goal epoch. Black line indicates circular-linear regression fit. C) Schematic of method for assessing goal-state phase precession. If a neuron exhibited a significant spike-phase spectral peak at relative frequency exceeding 1 (following multiple comparisons correction), and this effect was significantly stronger than that observed during trajectories to other goals, then this neuron was classified as exhibiting goal-state precession (*see Methods*). D) Example neurons exhibiting phase precession during navigation to specific goals. Left: Spike-phase spectra depicting frequency of neuronal spiking relative to ongoing LFP. Asterisk denotes spectral peaks that were significant and significantly different from other spike-phase spectra for other goals. Gray lines denote non-significant goals. Right: Spike-phase autocorelograms during navigation to each goal (significant goal epochs depicted in color). Text indicates the p-value for significance tests described in C).

We applied this method to the 448 non-spatially tuned neurons that were sufficiently active during each goal, and found a population of neurons exhibiting a significant pattern of faster-than-LFP rhythmic spiking during at least one goal (Fig. 6A), across a range of relative frequencies (Fig. 6B). Overall, 11% of these neurons (49/448) exhibited significant goal-state precession. We found at least one neuron exhibiting goal-state phase precession in 10/13 subjects. Of the neurons exhibiting goal-state precession, almost all did so for only one of the six goals (Fig. 6C). This effect was present at significant levels in anterior cingulate, orbitofrontal cortex, amygdala, and hippocampus, but not parahippocampal gyrus or entorhinal cortex (*p*’s 0.02, binomial test; Fig. 6D).

**Figure 6:**
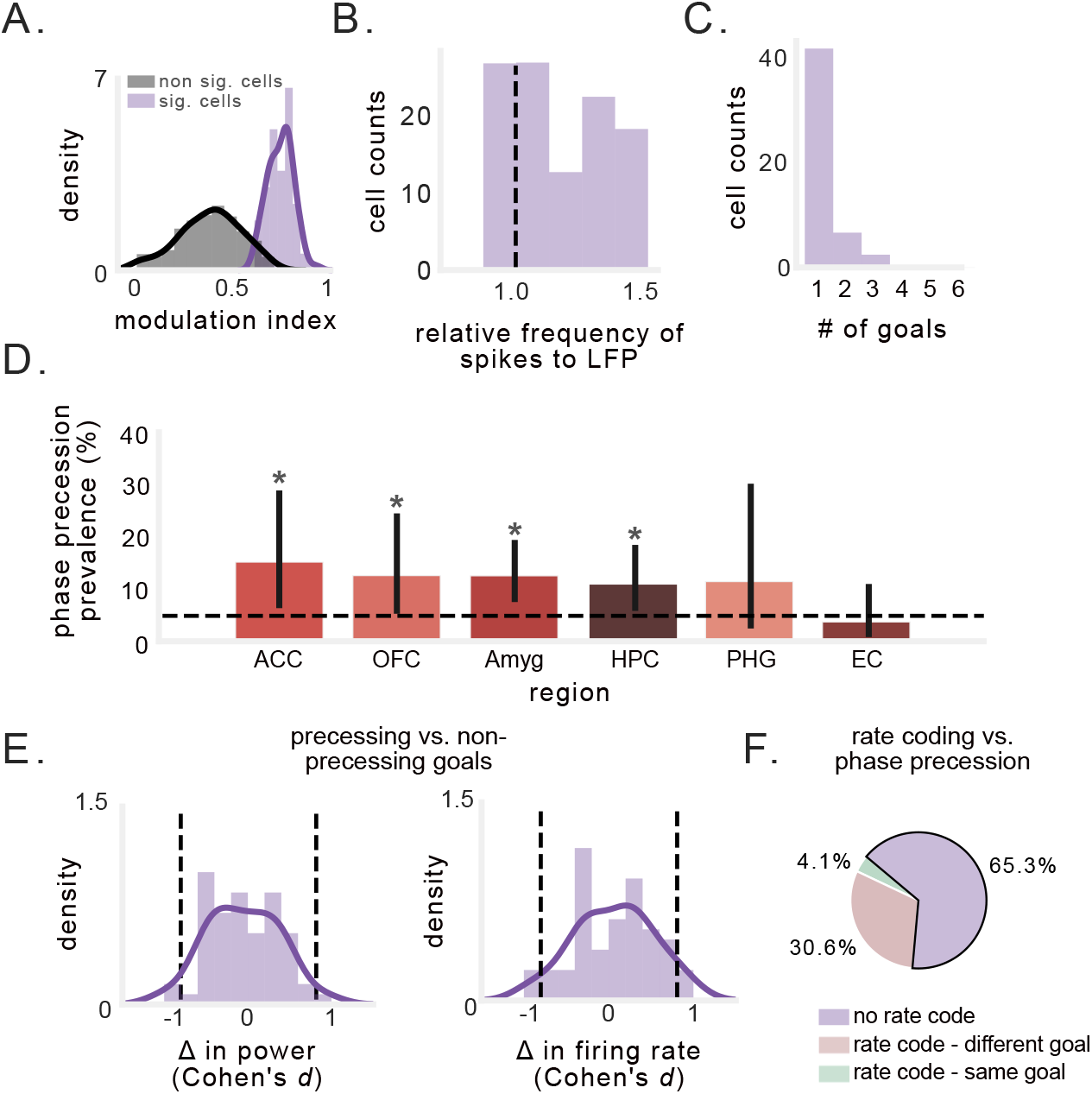
Prevalence and characteristics of goal-state phase precession in neurons that are not spatially tuned. A) Modulation index (MI) of spike-phase spectral peaks for significant vs. non-significant goals. B) Peak spike-phase PSD frequency for all goals for which a neuron exhibited a significant MI in the spike-phase spectra. Values to the right of the black line indicate that the neuronal frequency slightly exceeded the LFP frequency, indicating precession. C) Number of goals per neuron for which precession was observed. Most neurons exhibited precession during only one goal. D) Percentage of non-spatial cells in each region that exhibited goal-state phase precession. Asterisks indicate significant proportion of cells (*p*s< 0.02, binomial test). E) Distribution of Cohen’s d for the difference in 2–10 Hz power (left) and firing rate (right) between trajectories to goals showing precession vs. those that did not. Black dotted line indicate effect size of ±0.8. F) Analysis of overlap between goal-state phase precession and rate coding for goals.

We performed a series of control analyses that rules out the possibility that our observation of precession for specific goal states was confounded by between-goal differences in LFP power or neuronal firing rate (Fig. 6E). Indeed, neither example neuron in Figure 5 exhibited increased firing rates during goals that showed precession, suggesting that goal-state precession was independent of goal-specific firing rate increases^58^. Overall, only 17 of the 49 neurons that showed goal-state precession also showed increased goal-specific firing rate increases (*p*s< 0.05, one-way ANOVA), and only 2 of 17 of these neurons showed precession and a firing rate increase for the same goal^35^ (Fig. 6F). Next, we tested whether subject performance on different goals might be responsible for our results, i.e., whether precession might occur when subjects perform more efficient navigation. We measured subject’s performance on each goal (*see Methods*) and found no significant difference in navigational performance between goals that elicited precession and those that did not (Supplementary Fig. 9A, B). Because differences in theta power, firing rate, and behavior did not account for our results, our findings indicate that non-spatial phase precession selectively occurs during trajectories to specific goals and may also support the representation of non-spatial, sequential features of behavior.

## Discussion

The nature of the neural code is a fundamental question in neuroscience. Our findings show the first evidence that neurons in the human brain spike in rhythm with local network oscillations to represent spatial position and non-spatial events, in addition to the well-established code based on firing rate. Specifically, we demonstrate the existence of phase precession in humans during a virtual spatial memory task. We provide evidence for rodent-like spatial phase precession in human hippocampus and entorhinal cortex, in which spatially tuned neurons spike at earlier phases of theta (2–10 Hz) LFP oscillations as subjects moved through the putative place field center. We also provide evidence for the existence of non-spatial, goal-state phase precession which occurs transiently during trajectories to specific goals. These findings thus extend phase precession beyond rodents, and beyond spatial location, highlighting its potential as a more widespread neuronal mechanism for coordinating spike timing during behavior and cognition.

The spatial phase precession we observed bears important similarities to phase precession in rodents. We found spatial phase precession most predominantly in hippocampus and entorhinal cortex, where place and grid cells, respectively, are canonically found^2,59–61^. This suggests that the spatial phase precession we observed may be driven primarily by place- and grid-cells, as it is in rodents. One potential reason why phase precession has not previously been observed in humans is because human theta oscillations often appear at a slower and broader range of frequencies compared to those seen in rodents^43,62,63^. We specifically assessed phase precession relative to the broader range of theta frequency (2–10 Hz) fluctuations of the LFP, in line with the recent discoveries of phase precession in bats^55^ and marmosets^64^ — two animals with similarly heterogeneous theta oscillations. Our findings are also consistent with findings from rodents, who continue to show phase precession even when LFP theta power and theta-modulated spiking are reduced^25,65,66^. Furthermore, phase coding may not depend on regular, high amplitude rhythmicity in neural activity^51^. Instead, shifting LFP frequency can modulate spike-time intervals for synaptic plasticity without affecting the spike-phase^51^. Recent work in rodents has indeed demonstrated that the theta phase code is robust to changes in theta frequency, even as temporal lags between spikes are altered^56^. It is thus likely that the spatial phase precession we observed is analogous to that described in rodents, despite differences in theta range and rhythmicity.

Phase precession has predominantly been observed during place- or grid-cell spiking^67^. However, recent work has discovered the presence of phase precession relative to sound^33,34^, odor^34^, time in an episode^29–31^, task progression^27^, and REM sleep^29^. These findings highlight the potential generalizability of phase precession to non-spatial domains. In these instances, phase precession may enable the encoding of any successive stimuli or state, with the progression of phases binding a myriad of non-spatial sequences together for learning. By leveraging the idea that any variable may be encoded in spike phase if the frequency of spike rhythmicity exceeds the frequency of the local LFP oscillation^42,51^, we showed that phase precession also occurs with respect to behavioral states other than inhabiting a specific physical location—in this case, exclusively during trajectories to specific goals. The fact that this result is so specific, only showing up for a subset of goals for each neuron, might suggest an ensemble temporal code responsible for encoding all of the goals in the task^68,69^.

The goal-state phase precession we observed was largely independent of rate coding for goals, which has been described previously in human studies^58,60,70^. That the presence of rate and phase coding for goals is statistically independent is consistent with observations in rodents that showed that phase precession can appear for specific behavioral states even in the absence of concurrent firing-rate changes^35^. These findings support the theory that phase precession is used by the brain to signal behavioral states independent of firing-rate changes^71,72^. A challenge for future work is to understand the specific features of this phenomenon, such as the role of different phases within goal-state precession. One hypothesis is that goal-state precession may help track a person’s “episodic” position within a goal-seeking event. This would align with work in rodents showing phase-precession in “episode” or “time” cells when a rodent runs on a treadmill with a goal^30^ but not without a goal^73^, as well as evidence from human imaging showing that hippocampal and entorhinal cortex population activity correlates with distance to goal^74^. Furthermore, goal-state phase precession may relate to the phase precession observed in ventral striatum “ramp cells”^25^, and medial prefrontal cortex neurons in rodents^28^. The former exhibited precession as rodents approached reward locations, and the latter exhibited precession that was clearest when rodents approached the decision point in a maze^28^. Given that we found goal-state phase precession across various brain regions, including frontal cortex, these prior works further support the hypothesis that precession may represent “episodic” position within top-down behavioral states.

It is important to understand the prevalence of phase precession due to its likely theoretical relevance as neuronal mechanism for binding and compressing sequential events. In brief, phase precession organizes spiking at time intervals below the deactivation time constant of NMDA receptors, facilitating synaptic plasticity between neurons that encode events at behavioral time-scales^7,10,24,75–77^. Strengthening associations between consecutively active neurons may be a widely useful mechanism for learning associations. Our findings extend phase precession to the human brain and demonstrate that precession does not necessarily depend on physiological constraints, such as a stationary ~8 Hz theta oscillations^25,51,55,65,66^. Furthermore, our results show that a consistent difference between spiking and LFP frequency extends beyond place- or grid-field activity and may represent non-spatial mental sequences related to a memory task. This consistent spike–LFP frequency difference has been characterized by oscillatory-interference models as a function of spatial inputs, such as velocity, but may include non-spatial inputs if they increase spiking frequency above the network oscillation^78,79^. Therefore, our findings demonstrate the potential utility for phase precession in humans, across diverse brain regions, as a general, domain-free mechanism for temporal compression of specific experiences.

In summary, we have provided evidence for spatial phase precession in the human hippocampus and entorhinal cortex during virtual navigation and shown that it exhibits features similar to those seen in rodents. Further, we also demonstrated the existence of phase precession that is specific to trajectories to particular goals. These findings suggest that phase precession is a general mechanism for temporal coding in the human brain, despite the heterogeneity in theta rhythmicity in human MTL. Furthermore, the discovery of goal-state phase precession extends the potential for phase coding to be physiologically relevant for an array of experiential features, even when the neurons do not show concurrent firing rate changes for those features. Overall, our results suggest that phase precession is an important neural code across species and brain regions, not only for spatial cognition and memory but also for non-spatial features of experience.

## Methods

### Data recording and participants

The thirteen participants in our study were epilepsy patients who had Behnke–Fried microelectrodes^80^ (Ad-Tech Medical) surgically implanted in the course of clinical seizure mapping at the University of California, Los Angeles. The Medical Institutional Review Board at the University of California-Los Angeles approved this study (IRB 10–000973), and patients provided informed consent to participate in research. Microwire signals were recorded at 28–32 kHz, and we used Combinato for spike detection and sorting^81^. Manual sorting identified single-versus multi-unit activity versus noise on the basis of previously determined criteria^82,83^. The local field potential for each neuron was recorded from the local microelectrodes and was downsampled to 250 Hz for spectral analysis. For comparison with rodent data we used a publicly available dataset (CRCNS hc-2, hc-3)^41,42,84^.

### Task

This behavioral task is described in several previous studies^39,40,58,85^. Subjects first learned the navigational controls during a 4-delivery training session in a large, wide-open arena. After the practice session, subjects performed the main task, in which they were instructed to drive passengers to one of 6 goal stores in the virtual environment. Upon arrival, on-screen text displayed the name of the next randomly selected destination store. The task was self-paced in order to accommodate patient testing needs and therefore subjects performed at ceiling. The size of the virtual environment was 10 × 10 VR units, the width of the road was 2.5 VR units, and the obstructed area in the center of the road was 5 × 5 VR units. During navigation, subjects had a 60° field of view, a maximum forward speed of 1.25 VR units/s, a maximum backward speed was 0.5 VR units/s, and maximum angular velocity of 40°/s. To encourage subjects to take the shortest route to each destination, subjects received 50 points for each successful delivery and had one point deducted for each second that they spent navigating. Points were constantly displayed on-screen. Patients performed an average of 73 ± 11 deliveries in each session. To assess performance on this task, we measured subjects’ excess path length (EPL) on each trajectory, defined as ratio of the actual path length to the ideal path length. We computed ideal path length on each trial using the A-star search algorithm to identify the most computationally efficient path between goals in the environment^86^.

### Statistical analysis and software

All statistical analyses were carried out in Python, primarily using the SciPy^87^ and statsmodels^88^ libraries. For comparisons between two groups, we primarily utilized the Wilcoxon rank-sum test unless otherwise specified. For omnibus testing, we used ANOVAs, determining the p-value by comparing the real test-statistic to those from empirically derived null distributions generated by shuffling the true data. All figures were made using the Matplotlib^89^ and Seaborn libraries.

### Characterizing place-cell activity

To assess how neurnal activity related to the subject’s virtual spatial location, first, we binned the environment into a 10 × 10 spatial grid. We computed the firing rate map for each neuron by dividing the number of spikes by the amount of time spent in each spatial bin. We then used an ANOVA to assess whether the interaction of X and Y spatial bin (and thus 2D position) significantly modulated firing rate. To assess the significance of the ANOVA we circularly shuffled the firing rate and generated 500 surrogate test-statistics, and used this null distribution to determine the shuffle-corrected p-value of the ANOVA using the real data. These p-values were then FDR-corrected for multiple comparisons between the three movement types (CW, CCW, bi-directional).

Only neurons with critical statistics exceeding 99% of the shuffled data (*p* < 0.01) were considered to be spatially modulated. We considered spatially-modulated neurons to be analogous to place- and grid-cells, because the firing rate in at least one spatial bin deviated significantly from the others. We identified the spatial bin with the highest firing rate (analogous to the center of a place- or grid-field). We only included a spatial bin if the person passed through it at least 3 times, and occuped it for at least 4 seconds.

### Spectral analysis of LFP and spike time

To assess the prevalence and frequency of theta oscillations in the human and rodent LFP, we computed the continuous Morlet wavelet transform (wave number 6) at 20 logarithmically spaced frequencies between 1 and 32 Hz. Then, to identify theta-like oscillations, we utilized an iterative algorithm to subtract the aperiodic background and fit a Gaussian to putative peaks^90^. For this fitting procedure, we restricted the maximum number of peaks to 2, and the maximum peak width to 4 Hz. We only assessed the peak height (parameterized as the height of the Gaussian’s peak relative to the aperiodic background) and the peak frequency (parameterized as the center frequency at which the Gaussian reaches its peak) for the largest peak in the PSD. To assess the prevalence and frequency of theta oscillations in human and rodent spiking, we computed the autocorrelation of spike times, and performed a fast Fourier transform (FFT), yielding the PSD of the spike train.

### Phase estimation

We estimated the instantaneous phase of LFPs in the theta frequency range. Theta oscillations in human hippocampal formation are often bursty and non-stationary, and vary from low (2–5 Hz) to high (5–10 Hz) frequencies^43,62,63^ − similar patterns are observed in bats^91^ and non-human primates^92,93^. In order to analyze fluctuations in the LFP analogous to rodent theta, we estimated 2–10 Hz phase by identifying peaks, troughs, and midpoints in the lowpass-filtered LFP, and linearly interpolated between these points. This phase-interpolation method has been used previously to effectively estimate theta phase in bats^55^, as well as in rodents^94,95^. To ensure that phase estimates were not based on an unreliable low amplitude signal, we computed the instantaneous power of the LFP and discarded those time-points in which the power fell below a 25th percentile threshold^55^.

### Spatial phase precession

To identify phase precession in this dataset, for each spatially modulated cell we first identified every trajectory through the cell’s peak firing location. Following the methods used in some recent studies for measuring phase precession^55,96,97^, first for each such trajectory, we identified the spike closest to the center of the bin as the center spike (our reference point for the center of the bin on each trajectory)^55,96,97^. We limited our analysis to spikes in close spatial proximity to the center of the peak firing bin. To do so, we only analyzed the 11 closest spikes to the center of the peak firing bin. To ensure that these 11 spikes did not occur too distant from the peak firing bin, we set a diameter threshold of 40% of the environmental width, meaning that we did not analyze spikes that occurred further than 2 VR-units from the center of the peak firing bin. We re-ran our analyses while varying the inclusion criterion for the number of spikes (9, 11, & 13) and the radius (40% & 60%) and found that the parameters we selected did not significantly affect the proportion of cells exhibiting spatial phase precession (*χ*^2^ = 5.25, p = 0.5, chi-squared test).

We next tested for phase precession using circular statistics. Specifically, for each cell we measured the relation between spike phase and the subject’s position by computing the circular–linear correlation coefficient^47^. To assess statistical significance, we used a shuffling procedure. For each cell we computed a surrogate distribution for this correlation coefficient by assigning random phases to each spike from the distribution of all the spike phases for that neuron, and re-computing the correlation 500 times. This null distribution effectively scrambled the relationship between spike position and spike phase and controlled for any effect of spurious phase estimates. A circular-linear correlation was considered significant only if it exceeded the 95^th^ percentile of this surrogate distribution.

### Control analyses for spatial phase precession

We performed two control analyses for alternative explanations for the spatial phase precession we observed. The first analysis tested whether the peak firing bin, our analogue to the place-field center, was important to observing precession. To do so, we selected control locations for each cell and assessed the strength and prevalence in these control bins. Control bins were chosen as to not overlap with the peak firing bin (at least 30 % of the map width away) and also had to be traversed a minimum of 3 times with a minimum firing rate of 0.5 Hz. Furthermore, because we only analyzed the 11 spikes in the immediate vicinity of the peak firing bin, control bins matched the peak firing bin in sample size of spikes per trajectory, ensuring that effects were not confounded by firing rate differences.

Another possible alternative explanation for our findings is that the phase precession we observed here is actually encoding time to peak firing, independent of spatial position, with particular spike phases occurring at specific time-intervals within any epoch of elevated firing rate^98^. To control for this possibility, we identified epochs of elevated firing rate in the time domain without any information about position, which we refer to as “firing rate motifs”^98^. We identified the spike that occurred closest in time to the peak firing of each motif field, and used the 11 spikes in the immediate temporal vicinity (within 2 seconds before or after) to compute a circular-linear correlation between spike phase and spike time relative to the motif field peak, matching our spatial phase precession analysis.

### Non-spatial phase precession

To measure non-spatial phase precession without reference to place fields, we compared the spiking frequency of each neuron to the frequency of the local LFP, with relatively faster rhythmic spiking classified as phase precession^42,49^. However, detecting oscillations in spike times alone is difficult in humans (Supplementary Fig. 6) and bats^55^, potentially due to the transient, non-stationary nature of theta observed in these species^55,63^. Instead, we applied a method introduced by Mizuseki et al.^42^ which measures spiking frequency relative to the ongoing LFP. This is a particularly useful method when the ongoing LFP is non-stationary but may still be an important reference “clock” for neuronal spiking. To perform this procedure, we computed the autocorrelation histogram of each neuron based on the timescale determined by the phase of the reference LFP, rather than the conventional method of using absolute time. We computed this autocorrelation using 60° -bins with window-length of 4 cycles^55^. We then computed the Fourier transform of the autocorrelation histogram to yield the power spectral density (PSD) of the frequency of spiking relative to the LFP. Here, a peak relative frequency greater than 1.0 indicates that the cell is oscillating at a faster frequency than the reference LFP. We excluded neurons that exhibited both a peak near 1.0 as and significant phase-locking (*p* < 0.05, Rayleigh test) to ensure that we did not mistakenly identify phase-locked neurons^85^ as exhibiting phase precession. To measure the strength of this effect we measured the amplitude of the PSD, normalized by the total amplitude across all other relative frequencies^26^, which we refer to as the “modulation index” (MI) (Fig. 4A).

In order to ensure that our results did not arise from poor phase estimates due to low LFP amplitude, we discarded spikes that occurred during the lowest 25^th^ percentile of LFP power in the oscillation of interest^26,55^. In order to ensure that low spike counts did not confound our estimates we only analyzed cells with more than 100 valid spike-phase estimates for the autocorrelogram. We compared the modulation index to the null distribution of modulation indices for the peak frequency generated by circularly-shuffling the phases within each cycle of theta. This rigorous, within-cell shuffling ensured that cross-cycle dynamics (such as precession) were disrupted while maintaining slower and more rapid spiking dynamics^26,55^. The modulation index was considered significant if it exceeded the 95^th^ percentile of this surrogate distribution. Finally, we excluded cells that exhibited significant phase-locking during the entire session (Rayleigh test) in order to ensure that peaks close to 1.0 did not result from phase-locked spiking.

### Goal-state phase precession

To measure goal-state phase precession, we separately applied our analysis of non-spatial precession to spiking during each of the six goal trials. We only included neurons for which we observed at least 100 spikes per goal, to allow us to analyze non-spatial precession for each goal. We established two tests to characterize significant goal-state phase precession. First, just as we did with session-wide non-spatial precession, the magnitude of detected phase precession (as indicated by a peak in the spike-phase spectra exceeding 1.0) had to be greater than the 95^th^ percentile of the shuffled distribution. Because we conducted this test for all six goals separately, we used the False Discovery Rate procedure^99^ to correct the resulting p-values for multiple corrections across goals. If a goal exhibited significant non-spatial precession, we then compared the goal-specific modulation index to a surrogate distribution of modulation indices generated by selecting 500 random spike-trains from across the entire session. Each null spike-train was generated to match the number of spikes recorded during the significant goal to ensure that firing rate differences did not account for our results. A significant p-value indicated goal-state precession that was significantly stronger for the goal in question than for the session as a whole.

## Acknowledgements

We are grateful to the patients for participating in our study. This work was supported by the National Institute of Mental Health (R01-MH104606) and the National Science Foundation (to J.J.), and NSF Graduate Research Fellowship DGE 16-44869 (to S.E.Q.). We thank Kamran Diba, Sam McKenzie, Jonathan Miller, Andrew Watrous, and Melina Tsitsiklis for helpful comments and suggestions.

## Author Contributions

Conceptualization, J.J. and S.E.Q.; Methodology, J.J. and S.E.Q.; Investigation, I.F. and J.J.; Software, S.E.Q.; Formal Analysis, S.E.Q.; Writing – Original Draft, S.E.Q.; Writing – Review Editing, S.E.Q., J.J., and I.F.; Funding Acquisition, I.F., J.J., and S.E.Q.; Resources, I.F. and J.J..; Visualization, S.E.Q. and J.J.; Supervision, J.J.

## Declaration of Interests

The authors declare no competing interests.

---

**Supplemental Figure 1:**
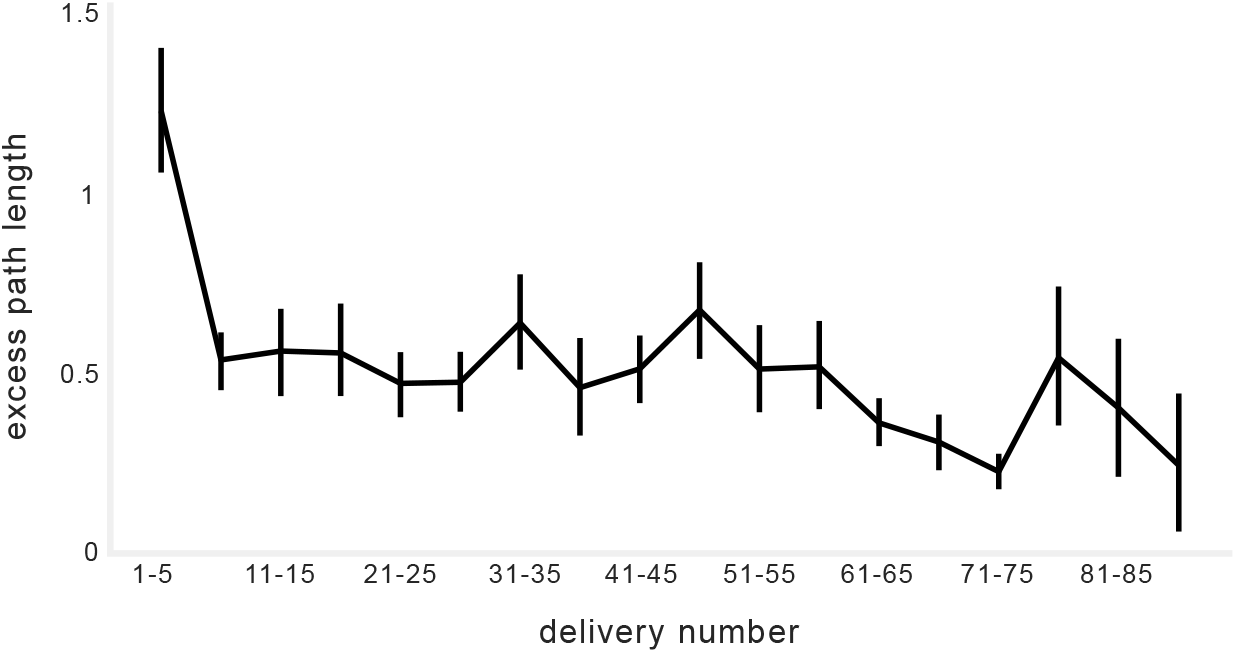
Behavioral performance in the task. Plot indicates the mean excess path length across all sessions as a function of trial number.

**Supplemental Figure 2:**
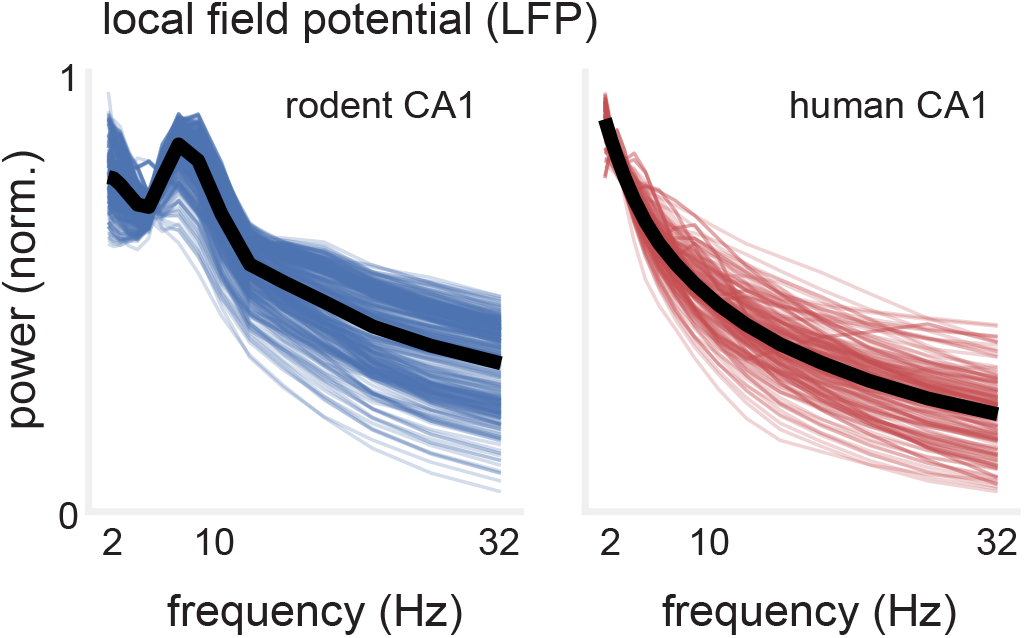
Differences in rodent and human hippocampal theta oscillations. PSD of hippocampal LFPs recorded in navigating rodents (blue) and humans (red). Black line denotes average across channels. Rodent hippocampal LFP shows a clear peak in the 5-10 Hz range in almost all channels, while the human LFP does not.

**Supplemental Figure 3:**
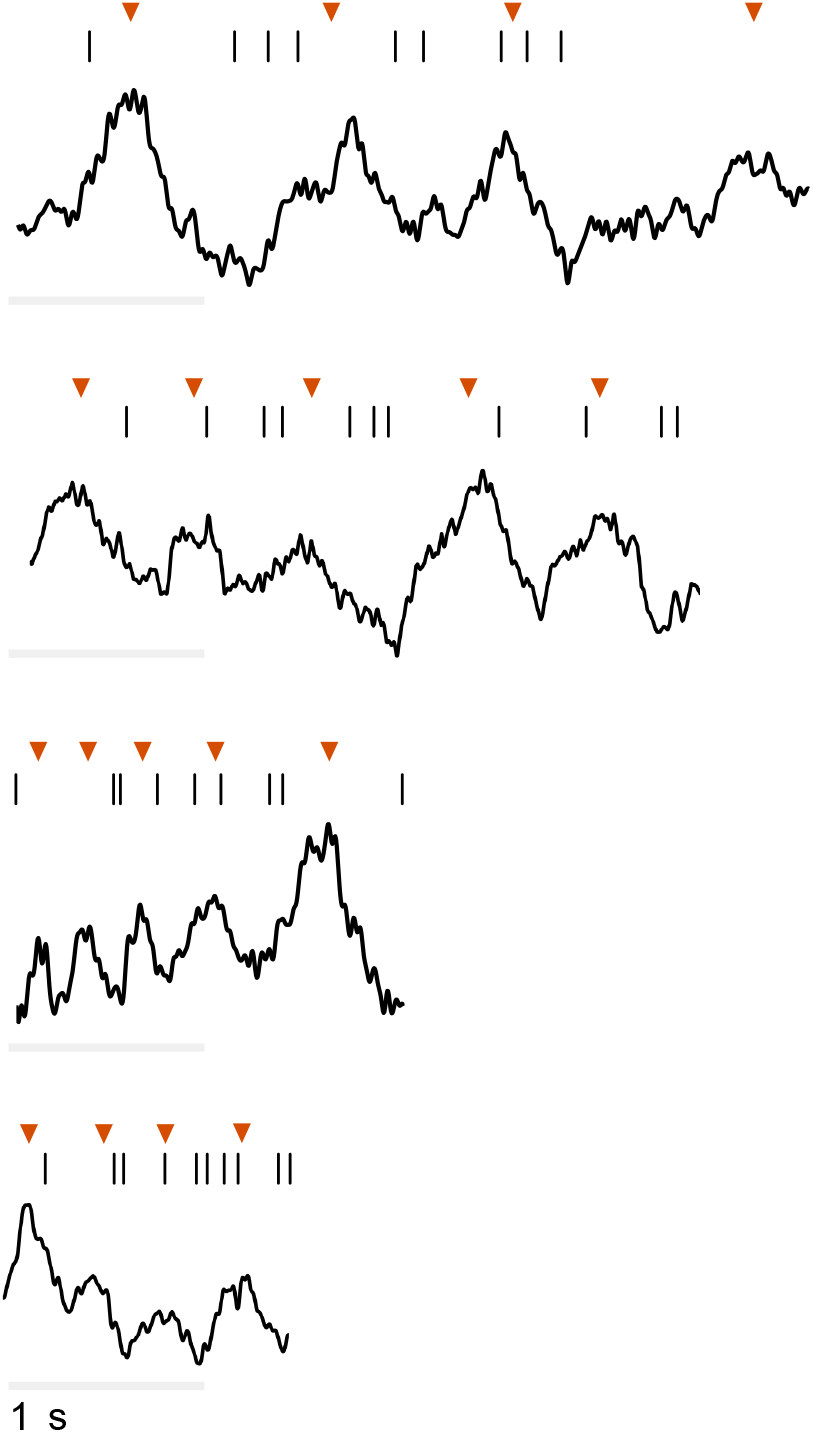
Examples of spatial phase precession during individual passes through a field. Spike times and 1–30-Hz filtered LFP data during individual passes through peak firing bins for four neurons that exhibited significant spatial phase precession. Red arrows denote peaks of individual theta cycles.

**Supplemental Figure 4:**
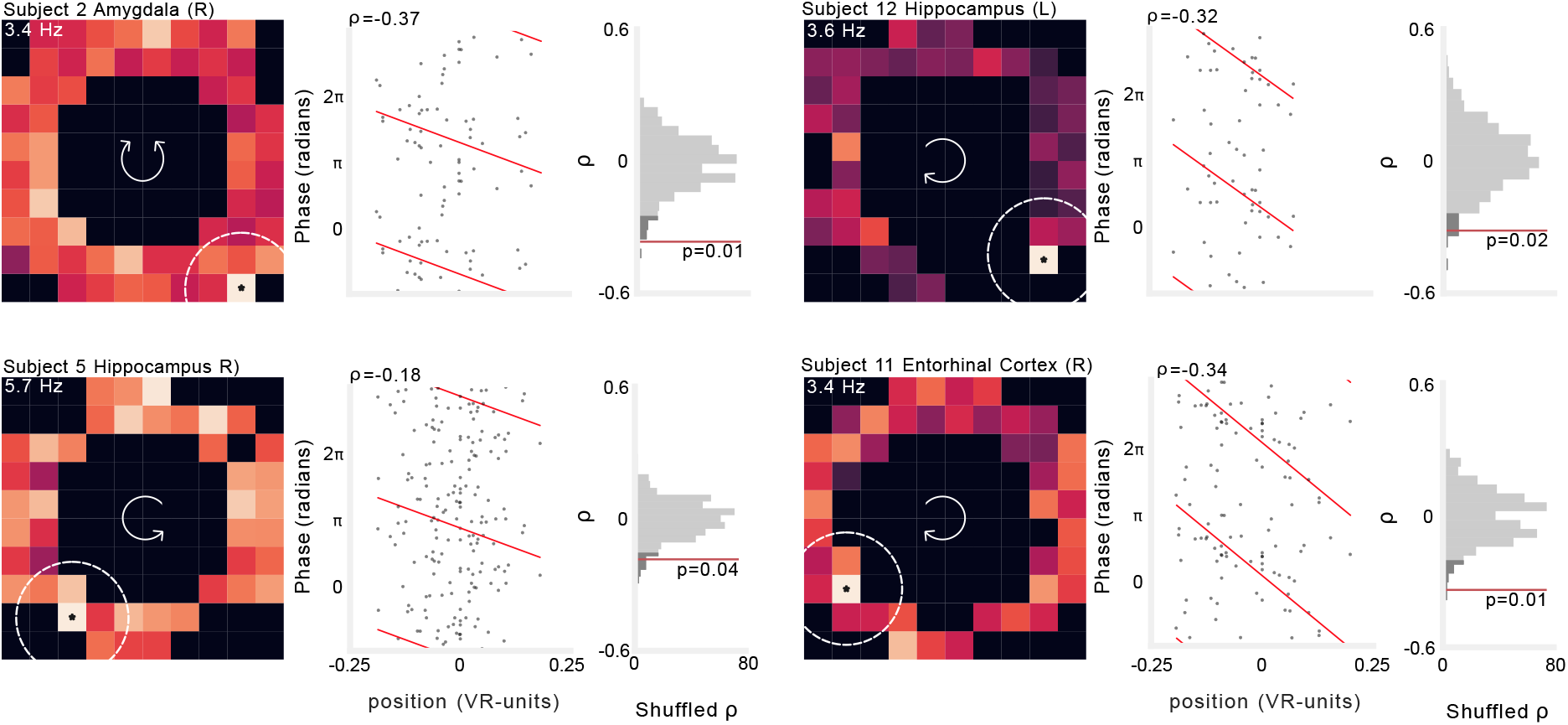
Additional examples of spatial phase precession. The activity of four neurons that show significant spatial phase precession. Left: firing rate heat map. Brighter colors denote higher firing rates. Text label indicates the color scale for the plot with the mean firing rate of the peak firing bin, which is noted with an asterisk. Dotted lines indicate maximum radius around field in which spiking was assessed. Arrows in the center of the heatmap indicate the movement direction for which this plot was computed. Middle: spike phase as a function of location relative to the field center. Spike phases are duplicated vertically to enable visualization of circular–linear regression (red). Text indicates circular-linear regression coefficient (rho). Right: surrogate distribution of circular-linear regression rho-values generated by permutation of spike phases. Red line indicates value of real data. Dark gray shading indicates 95^th^ percentile of surrogate distribution.

**Supplemental Figure 5:**
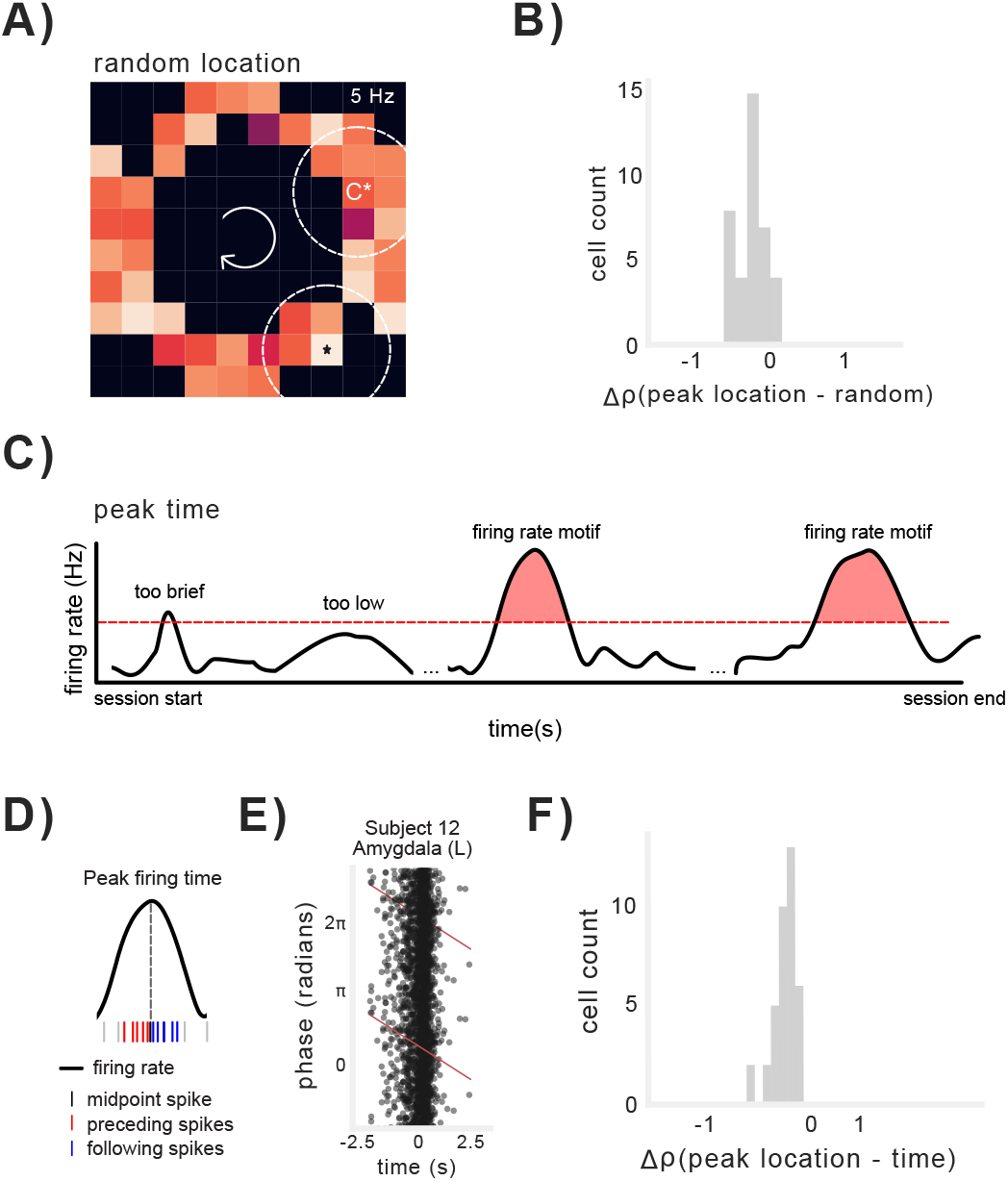
Location- and time-control analyses for spatial phase precession. A) Example of alternate location selected to test whether peak firing bins exhibited significantly greater phase precession than randomly selected locations. B) Distribution of differences in circular-linear correlation coefficients for precession observed in peak firing bin vs. randomly selected location. C) Schematic of method for identifying elevated firing rate. Firing rate had to exceed a firing rate threshold of 1.5 Hz for at least 250 ms in order to be classified as a firing rate “motif”. D) Schematic of of method for time-based phase precession within motifs of elevated firing rate. E) Example neuron exhibiting significant phase precession relative to elapsed time within a firing rate motif. F) Distribution of differences in circular-linear correlation coefficients for precession observed in peak firing bin vs. time in firing motifs.

**Supplemental Figure 6:**
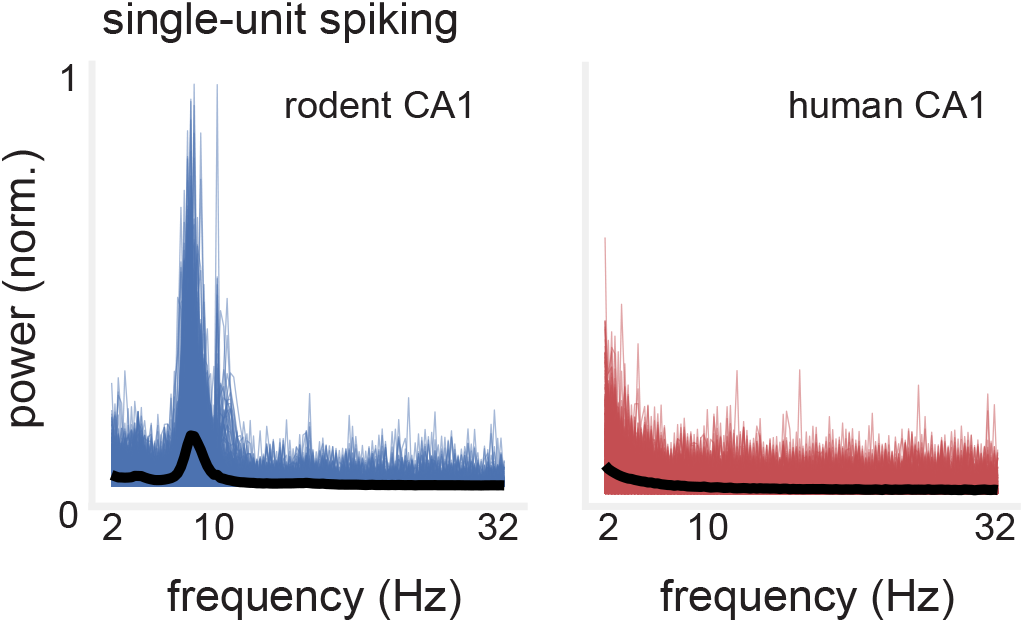
Differences in rodent and human hippocampal neuronal spiking. Power spectral density from single-unit discharge from rodent (blue) and human (red) hippocampus. Solid black line indicates average across neurons. Rodent spiking shows clear theta modulation of spike timing while human spiking does not.

**Supplemental Figure 7:**
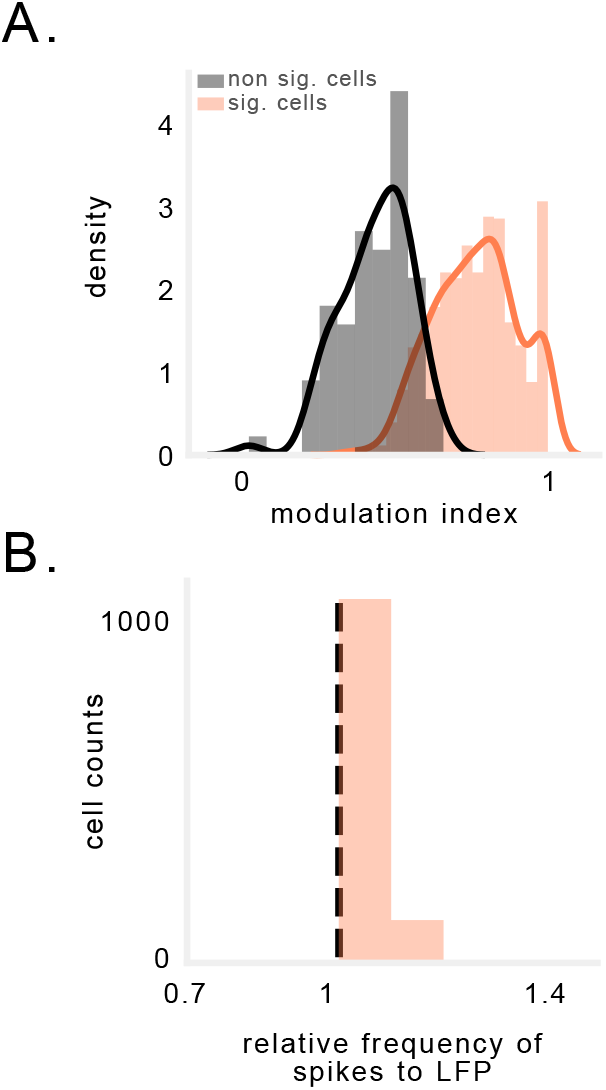
Phase precession in rodent CA1 without reference to position. A) Modulation index (MI) of spike-phase spectral peaks for significant vs. non-significant neurons recorded in rodent CA1. D) Distribution of relative frequencies for neurons exhibiting significant MI in the spike-phase spectra. Values to the right of the black line indicate that the neuronal frequency slightly exceeded the LFP frequency.

**Supplemental Figure 8:**
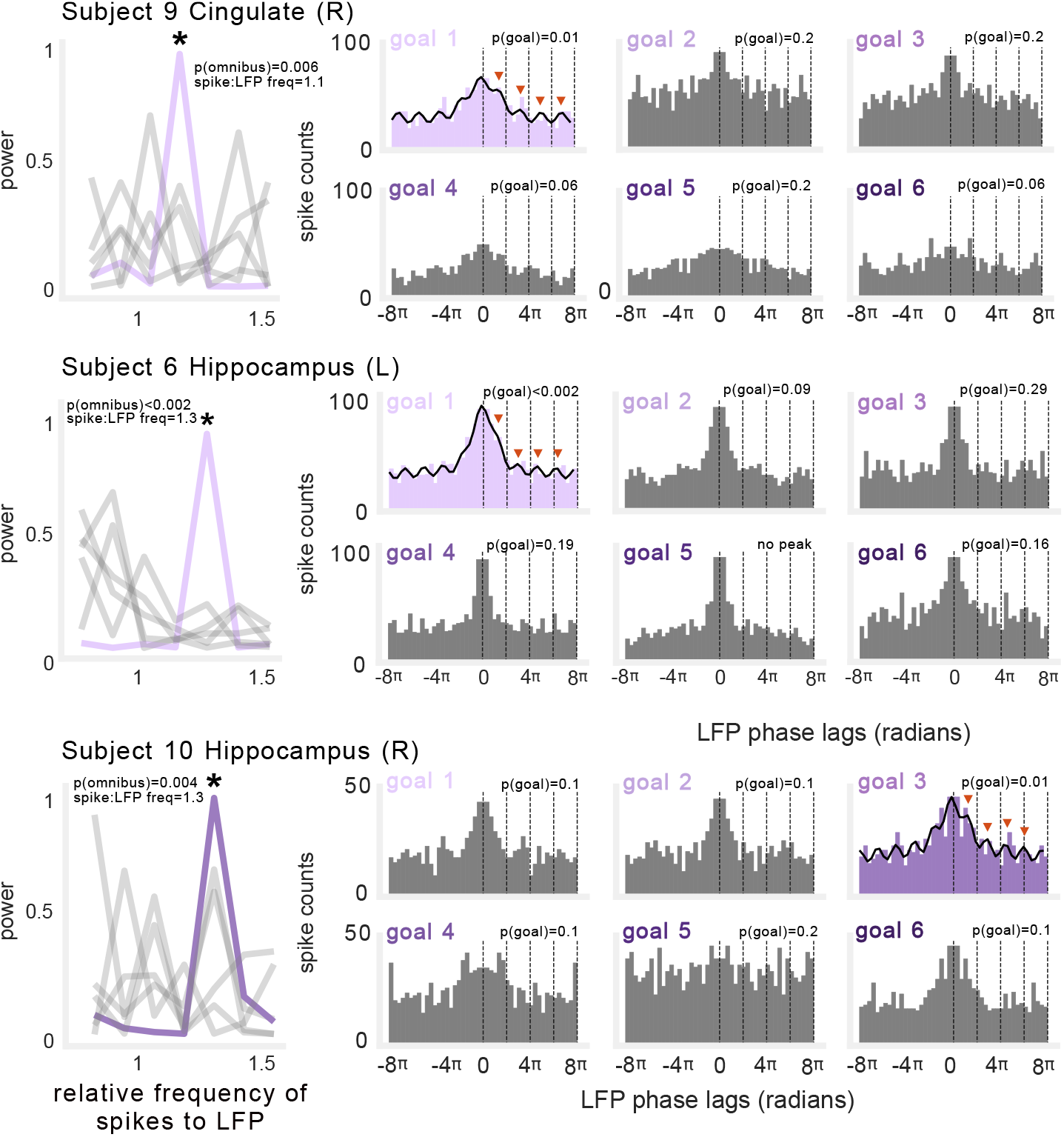
Additional examples of goal-specific phase precession. Three example neurons exhibiting phase precession during navigation to specific goals. Left: Power spectral density depicting frequency of neuronal spiking relative to ongoing LFP. Asterisk denotes peaks that were significant and significantly different from other goals. Gray lines denote spike-phase spectra for non-significant goals. Right: Spike-phase autocorelograms during navigation to each goal (significant goal epochs depicted in color). Text indicates the p-value for both significance tests described in Figure 5C), and relative frequency of spiking to LFP. Black line indicates fit of decaying-oscillation function added to significant examples shows oscillation in spike-phase autocorrelogram.

**Supplemental Figure 9:**
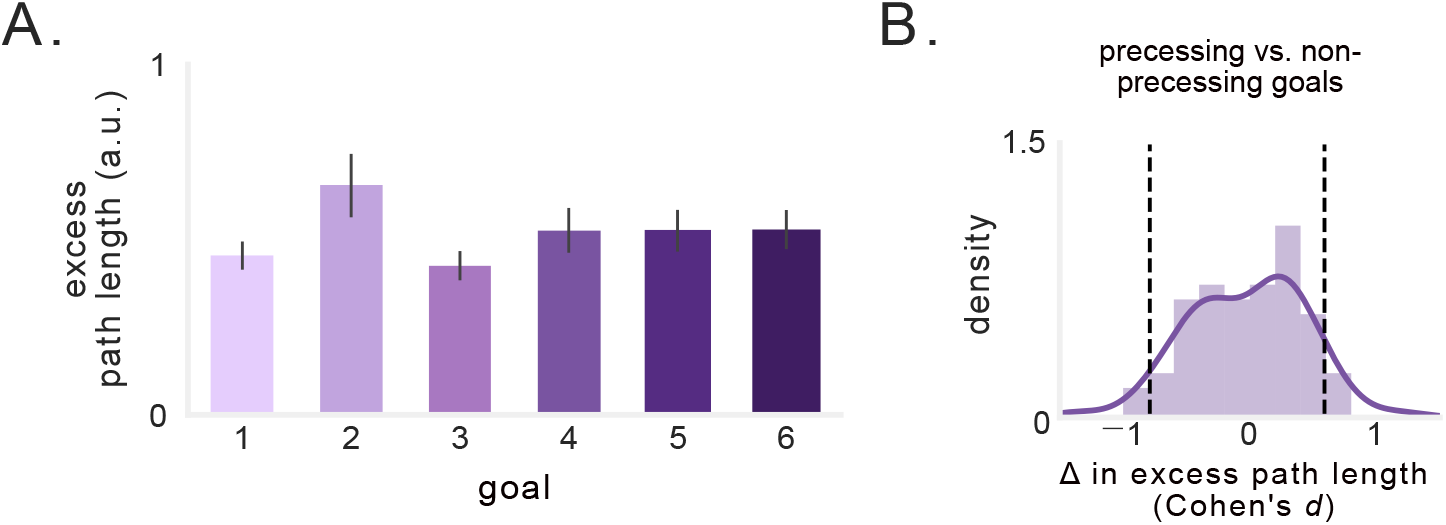
Goal-specific phase precession is not a function of a differences in navigation performance. A) Excess path length as a function of goal. B) Distribution of Cohen’s d comparing excess path length during trajectories to goals that showed precession vs. those that did not. Black dotted line indicates effect size of ±0.8.

## Notes

### Competing Interest Statement

The authors have declared no competing interest.

